# Mutualism provides the basis for biodiversity in eco-evolutionary community assembly

**DOI:** 10.1101/2024.02.02.578708

**Authors:** Gui Araujo, Miguel Lurgi

## Abstract

Unveiling the ecological and evolutionary mechanisms underpinning the assembly of stable and complex ecosystems is a main focus of community ecology. Ecological theory predicts the necessity of structural constraints on the network of species interactions to allow for growth of complexity in assembling multispecies communities. A promising research avenue is the search for an understanding of how the coexistence of diverse species interaction types could influence the development of complexity and how an ideal composition could arise in nature. We propose an ecological model with mixed interaction types incorporating evolutionary assembly by speciation. This framework allows to investigate the eco-evolutionary assembly on complex species interaction networks with multiple interaction types and its consequences for ecosystem stability. Our results show that highly mutualistic communities are conducive of complexity and promote the emergence of consumer-resource interactions. Furthermore, we show that an evolutionary process is required to produce such condition. Moreover, this evolutionary assembly generates a diversity of outcomes and promotes two distinct types of complexity depending on speciation constraints. Assembled communities are thus either larger (more species) or more connected, in agreement with patterns previously observed in microbial communities. Our results produce invaluable theoretical insight into the mechanisms behind the emergence of ecological complexity and into the roles of mutualism and speciation on community formation.

## 1 Introduction

Understanding the mechanisms responsible for the generation and maintenance of diversity in complex natural communities is a central focus of ecological research [1, 2]. Both evolutionary mechanisms generating diversity (e.g. speciation) as well as ecological assembly processes driven by biotic and abiotic conditions (e.g. environment, species interactions), have been shown to be capable of generating the complexity observed in these communities[2–4]. The challenge remains, however, of understanding the relative importance of different mechanisms acting together to foster biodiversity while at the same time maintaining the stability of assembled communities[5, 6]. A better understanding of the mechanisms, and their interplay, that generate, maintain, and disrupt biodiversity, is fundamental to our conservation efforts to preserve it[7, 8].

During the assembly process, communities grow by the addition, either ecological (e.g. via immigration / invasion) or evolutionary (e.g. via speciation), of new species. These new species must survive, establish, and grow in the ecological context in order to become part of the community. Changes prompted by this addition can disrupt the structure of the recipient community, potentially resulting in the extinction of resident species, a process mediated by the ecological interactions present in the community. Nonetheless, it is through these processes of assembly that communities grow, providing a platform for the emergence of community-level patterns while challenging the stability of the community.

Five decades ago, May[9] proved that more complex ecological networks (in terms of species richness and connectivity) are less likely to persist than relatively simple ones. This seminal finding initiated a several-decades long quest for the organisational patterns of natural communities that allow them to grow in complexity, apparently without compromising persistence and stability. May’s result referred to randomly connected communities (i.e. not subjected to a deterministic assembly process), with an arbitrarily large number of species and connections. We now know that specific network structures can modify this complexity-stability relationship[10], and that in empirical systems a large number of species often translates into low connectivity[2]. The next challenge is to identify the mechanisms behind the emergence of these structural patterns and the diversity-connectivity relationships.

From an ecological point of view, both deterministic (e.g. biotic interactions) and stochastic (e.g. colonisation) processes influence the outcome of community assembly. Priority effects, given by stochastic colonisation, can have profound consequences for the final outcome of community assembly[11, 12]. Once species manage to colonise a given habitat, ecological interactions take over, strongly influencing the resulting community. In this sense, the relative composition of different interaction types has been shown in theoretical studies to be a strong determinant of community assembly and stability[13, 14]. However, contrasting hypotheses regarding the role of interaction types on community stability exist, especially about the role of mutualistic interactions for community persistence[15, 16]. Some theoretical studies on community assembly by repeated invasion of new species suggest that stability is achieved with a mixture of types, with a majority of mutualisms[13, 17, 18]. But the opposite has also been shown, with some further results showing that mutualism promotes instability while competition or antagonism promotes stability[10, 19]. A potential route towards consensus across these contrasting results is to consider the idiosyncratic effects of mutualisms depending on their functioning and distribution within the network of species interactions. The role played by mutualisms in the community and their interplay with other interaction types may be a key determinant of the assembly process. Additionally, studies of stability tend to focus on species richness and composition, describing the effects of perturbations in terms of subsequent extinctions. However, interaction types might reveal a more straightforward role in community structure if considered from a different point of view, such as their direct relationship with complexity.

From an evolutionary perspective, a major contributor to community assembly is speciation. By generating diversity from existing species and traits, the process of speciation, and the corresponding selection that ensues, is capable of generating biodiversity patterns similar to those observed in real ecological networks[20, 21]. Investigating community assembly by considering speciation can thus help us understand the emergence of complexity in ecological communities. Further, by comparing evolutionary assembly by speciation with ecological assembly by repeated invasions we can better understand the contribution of evolutionary processes to the assembly of ecological communities[22]. So far the main focus of evolutionary assembly models has been on the study of either food webs or mutualistic networks independently[20– 25]. The interplay of mutualism, competition, and consumer-resource interaction types together in evolutionary models of assembly as a mechanism for the generation of complexity has not so far been considered.

Here, we fill this gap by investigating the evolutionary assembly of complex ecological networks comprising different types of ecological interactions using a theoretical model. We consider the interplay between mutualistic, consumer-resource, and competitive interactions in an eco-evolutionary setting. We model the evolutionary assembly of communities using a dynamical model determining the abundance of species (ecology) as they are introduced into the community via a process of speciation (evolution). Interactions dictate the influences species have on one another and can lead to their dynamical extinction. The outcome is then a stable structure in the species interaction networks shaped by this process of ecological selection. We assess the effects of assembly via evolutionary speciation by contrasting the eco-evolutionary model with a model of ecological invasion. The influence of mutualisms on assembly are assessed by comparing scenarios with and without mutualistic interactions. We compare eco-evolutionary trajectories with purely ecological ones (grounded on species invasions) to disentangle the effects of ecology and evolution and to determine which outcomes are feasible from naturally selected parameters. We quantify structural properties of the network of ecological interactions and relate their changes along the assembly process to changes in species richness and complexity. This allows us to unveil mechanisms for species persistence and community stability and how they are brought about by evolutionary changes.

We find that mutualism and speciation play a key role in the increase of complexity and species diversification. However, they are not necessary for an increase in species richness, which can happen at the cost of low network connectivity in purely ecological scenarios in the absence of mutualisms. Our results show that a high proportion of mutualism allows the growth of complexity and breeds the generation of consumer-resource interactions. Evolutionary speciation is necessary to enable the emergence of this outcome. Evolutionary assembly also generates diversity of outcomes and produces distinct types of complex mutualistic communities. This is modulated by the magnitude of change between parent and offspring species in speciation events. Our results match empirical patterns of richness-connectance relations observed in microbial communities and suggest an evolutionary mechanism for their occurrence.

## 2 Results

### 2.1 Mutualisms promote complexity

A model of ecological assembly via repeated invasions selects against competitive (*−/ −*) interactions while favouring mutualistic (+*/*+) ones. However, this selection is weak (Fig. 1). A high proportion of mutualisms only happens when this type of interaction is forced by the model’s parameters. Across four different invasion scenarios (enforcing mutualisms to high fractions, without mutualisms, enforced balanced interaction types, or random), species richness is generally low. However an increase in richness is observed when mutualisms are not present, and an even larger increase is observed with a high proportion of mutualisms (Fig. 2A). This increase in richness coincides with a decrease in connectance and complexity for the model without mutualisms (compared to the random scenario), but with an increase in complexity when high fractions of mutualisms are enforced (Fig. 2B-C). Species abundances are also influenced by mutualisms, with forced mutualistic communities capable of attaining high biomass both at the community and the species level. In the absence of mutualisms, on the other hand, a decrease in the abundance per species is observed (Fig. 2E-F).

**Fig. 1.**
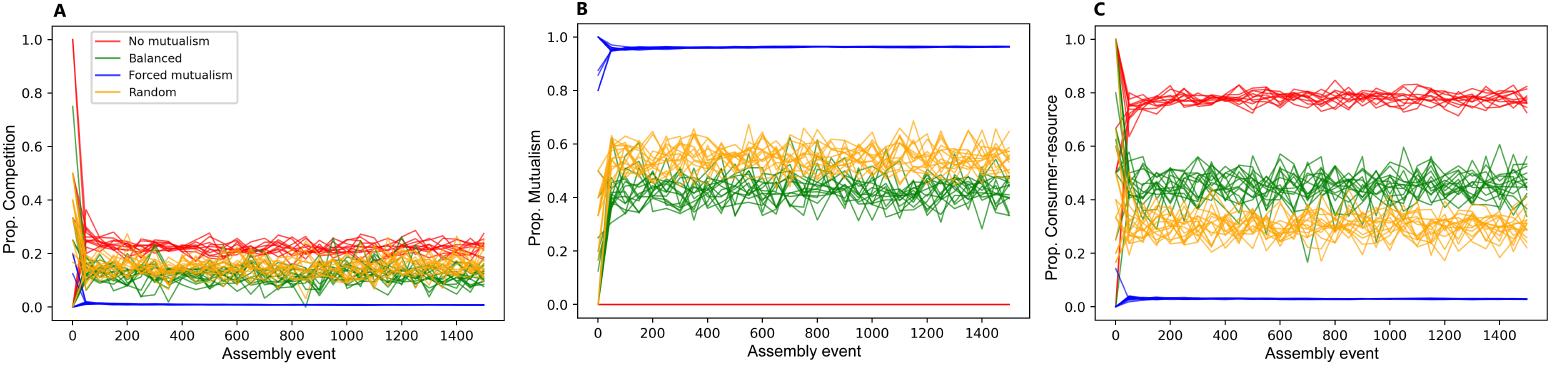
Assembly by invasion: interaction types. Composition of interaction types during assembly for each variation of the ecological assembly model via repeated invasions: without mutualism, fixed balanced proportion of interactions, high mutualistic fractions, and randomly chosen proportions of interaction types. All cases present very stable and convergent assembly trajectories, without a variation of outcomes. **(A)** Competitive interactions are selected against and decrease in proportion in all scenarios. **(B)** Mutualism is selected for in all scenarios and presents a slightly higher proportion when there is freedom in interaction type selection in comparison with fixed balanced proportions (orange and green). **(C)** Consumer-resource interactions present more neutrality in selection and give space to more mutualism in the random case, reflecting the favouring of mutualistic interactions.

**Fig. 2.**
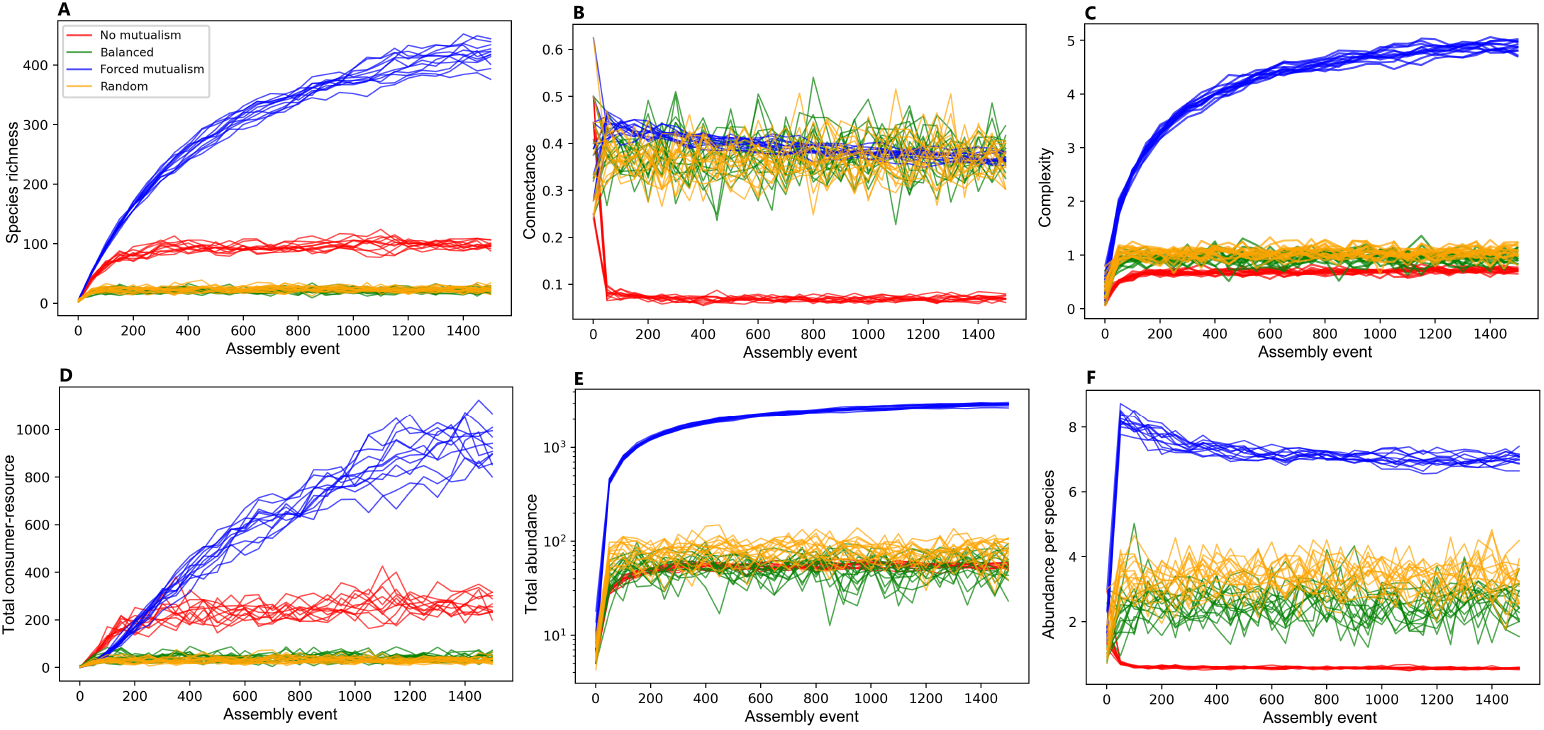
Assembly by invasion: community complexity. Trajecories of the variables’ histories across assembly events, during which new species are added to the community at equilibrium, for each variant of the invasion models. **(A)** Without mutualism, richness increases. However, with forced high mutualistic fractions, it increases considerably more. **(B)** Without mutualism, the increased richness comes with the cost of less connectivity between species (measured as network connectance (*C* = *L/S*^2^), but the same is not observed for high mutualism. **(C)** Therefore, only high mutualism allows an increase in complexity 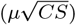, but this scenario is not selected naturally with assembly by invasion (from the random model). It can only be forced by hand. **(D)** Furthermore, even consumer-resource interactions become more numerous with high mutualism, despite their much lesser proportion. **(E)-(F)** High mutualism also allows for more abundance both at the community and species levels, while the growth in richness without mutualism occurs at the expense of abundance per species.

These findings show that only highly mutualistic networks can promote complexity. Furthermore, despite the high fraction of mutualistic interactions, consumer-resource (+*/−*) ones also become more prevalent (Fig. 2D). Mutualisms thus promote the maintenance of other interaction types.

To investigate the routes through which mutualisms influence complexity, we introduced mutualistic interactions to communities assembled without them. Interestingly, this prompted a decrease in species richness. The removal of mutualistic interactions from a balanced model, on the other hand, prompts an increase in species richness (Fig. 3A-B). Despite destabilising the community and the resulting drop in richness, the introduction of mutualisms increases network complexity by increasing connectance (Fig. 3C).

**Fig. 3.**
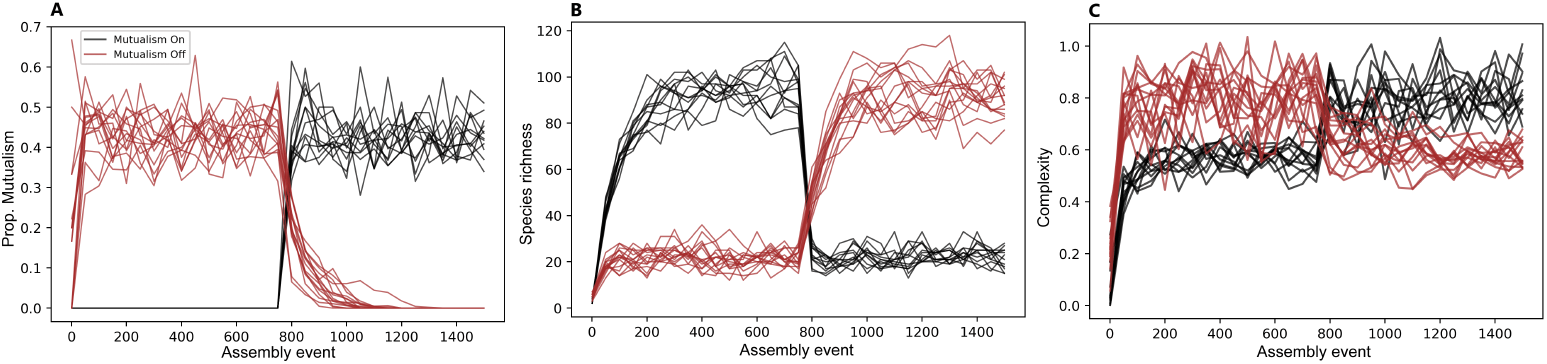
Mutualism is about complexity, not richness. Consequence of switching mutualisms on in a model without them and switching them off in a balanced model with mutualisms. **(A)** By switching off, mutualisms disappear completely. By switching on, the community restores its usual composition. **(B)** The introduction of mutualisms causes a decrease in species richness, while its removal causes an increase. **(C)** Despite that, the change in complexity is the opposite.

These findings suggest that mutualism decreases richness when in low proportions and increases it dramatically in large proportions but, regardless of the composition, it increases complexity. Our results show that a high proportion of mutualism begets complexity, but a caveat of invasion models is that they are not capable of facilitating the emergence of predominantly mutualistic communities, even when proportions of interaction types are freely allowed to change (random invasion model). This configuration of interaction types has to be enforced. It is also notable how convergent all invasion models are, producing samples of communities with almost no variety of possible outcomes.

### 2.2 Evolution is a key driver of diversity and complexity

The evolutionary speciation model naturally selects for communities (i.e. networks) with high proportions of mutualistic interactions, high abundance per species, and high species richness. This is observed particularly when the magnitude of change in evolutionary events is constrained (Δ = 5), resulting in a much higher species richness compared to the forced mutualisms invasion scenario (Fig. 4). Compared to the random invasion model (the purely ecological null counterpart of the evolutionary speciation model), both constrained (Δ = 5) and less constrained (Δ = 30) evolutionary scenarios are capable of generating larger and more complex communities (Fig. 4A-C).

**Fig. 4.**
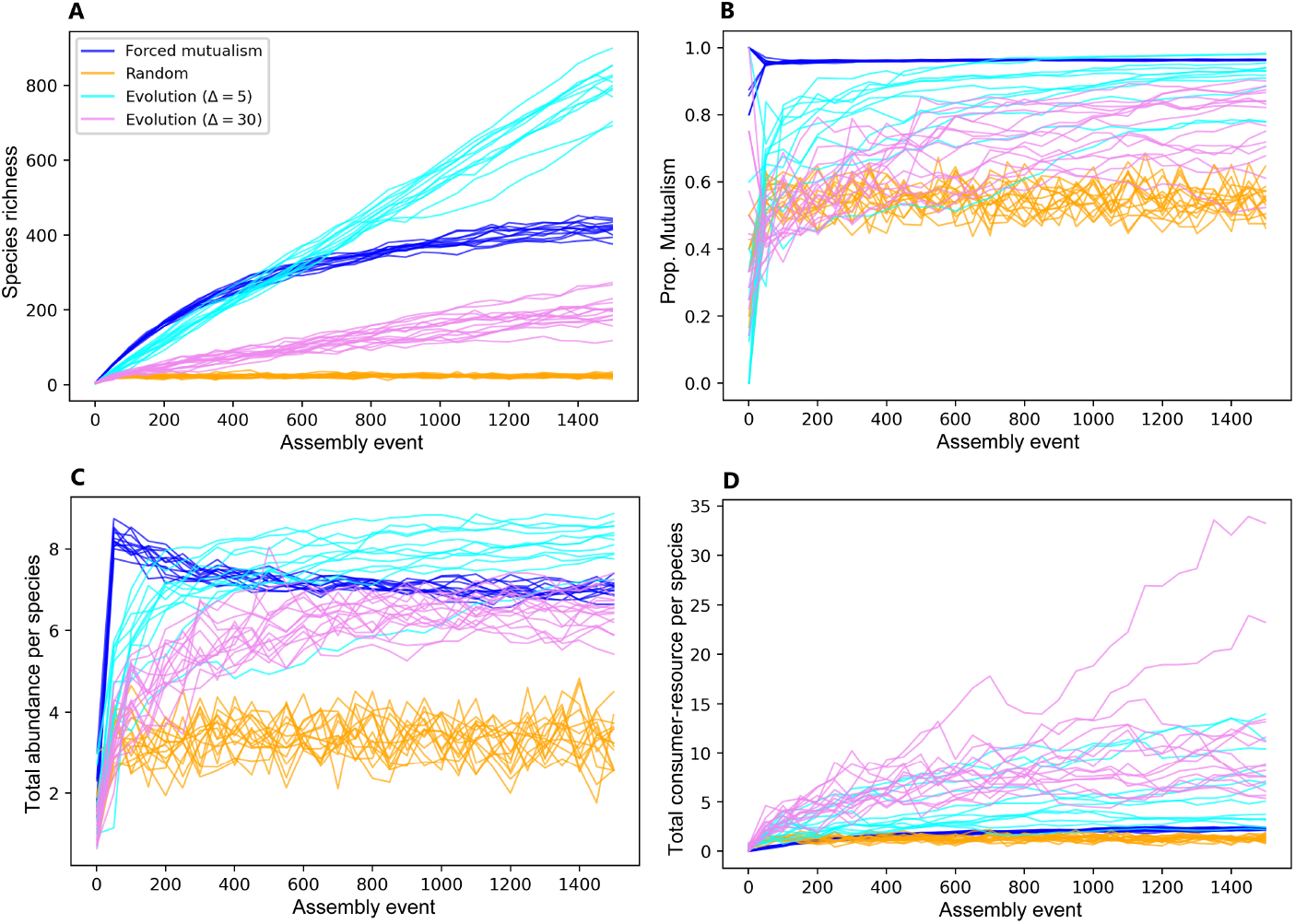
Assembly by evolutionary speciation generates diversity. Communities assembled via evolution exhibit a larger variation of outcomes. Evolution results in the highest species richness **(A)** by effectively selecting a high proportion of mutualisms **(B)**. A more controlled evolution (Δ = 5) is more effective. It also produces species that are more abundant on average **(C)**. The amount of consumer-resource interactions per species is also larger, with a less controlled evolution (Δ = 30) being more effective at this **(D)**. Therefore, evolution also enables the emergence of more abundant consumer-resource relations.

The model with less constrained evolution (Δ = 30) is less able to promote species richness than its more constrained counterpart (Δ = 5). However, it is able to harbour more consumer-resource interactions per species, even though both evolutionary models exhibit higher levels of consumer-resource interactions when compared to the invasion assembly models (Fig. 4D). With a higher richness and a higher number of consumer-resource interactions per species, the high proportion of mutualistic interactions from evolved communities acts as a stage for the establishment of food webs.

Intriguingly, the evolutionary assembly of communities is capable of generating a larger variety of outcomes across sample communities compared to any of the invasion models. This diversity of outcomes represents an additional layer of complexity being generated when considering a collection of communities produced by an evolutionary assembly process. Evolutionarily assembled communities present high levels of complexity distributed across a wide range of values (Fig. 5A). Comparing between two levels of evolutionary control, we observe the emergence of two distinct types of complex communities. Constrained evolution produces higher species richness and lower connectance, compared to the less constrained evolutionary scenario (Fig. 5B). In terms of the strength of ecological interactions, we observe a convergence across both ecological and evolutionary scenarios (Fig. 5C), demonstrating that the differences in complexity are only due to differences in richness and connectance across different community types.

**Fig. 5.**
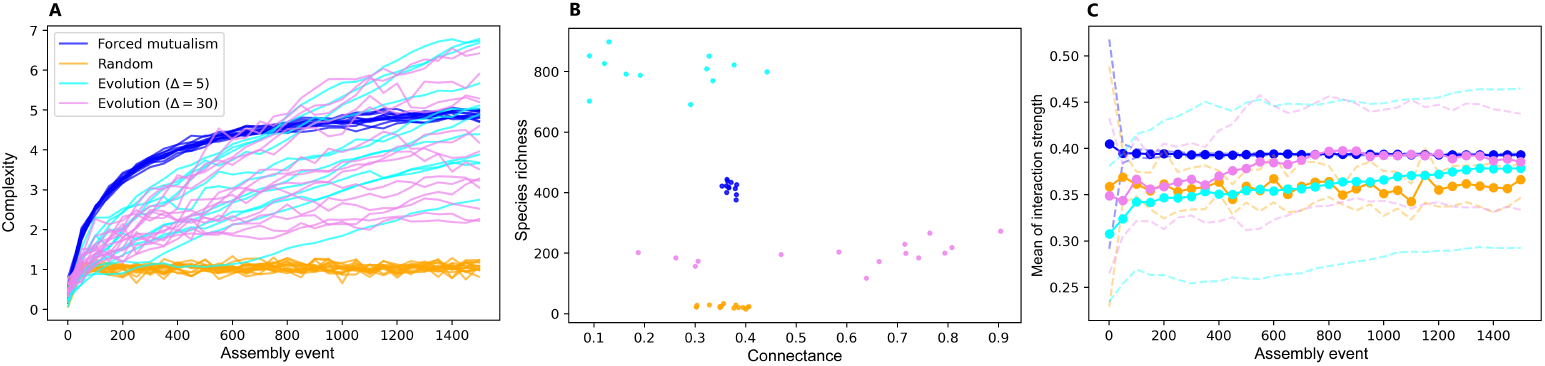
Evolutionary selection favors complexity. **(A)** Evolution naturally reaches levels of complexity comparable to the high mutualism variant of the invasion model (in which mutualistic interactions are enforced) and is able to reach a wide range of complexity values. **(B)-(C)** With very similar profiles of complexity, a more controlled evolution (Δ = 5) selects for more richness, while a less controlled evolution (Δ = 30) selects for more connectivity and higher variance of interaction strengths. **(C)** Shows average trajectories with standard deviation boundaries, dotted at every 50 assembly events.

### 2.3 Networks are structured by mutualistic interactions

Highly mutualistic communities, both ecologically and evolutionarily assembled, feature a higher diversity of species’ connectivities (i.e. degree entropy) than expected by chance (Fig. 6A). This suggests that mutualisms tend to spread in diverse ways across assembled networks, with a more even distribution of degrees and without dominance of particular degree values. This effect is stronger with more controlled evolution (Δ = 5), which also presents smaller connectance. Our results also suggest that there is a selection pressure for higher modularity, as communities exposed to strong selection (random invasion and evolutionary assembly) feature a higher modularity than expected by chance (Fig. 7B). Again, this effect is stronger for more constrained evolution, which makes intuitive sense, since complex communities with a lower connectance can be expected to be more effectively separated into modules and selected for modularity when compared to highly connected communities.

**Fig. 6.**
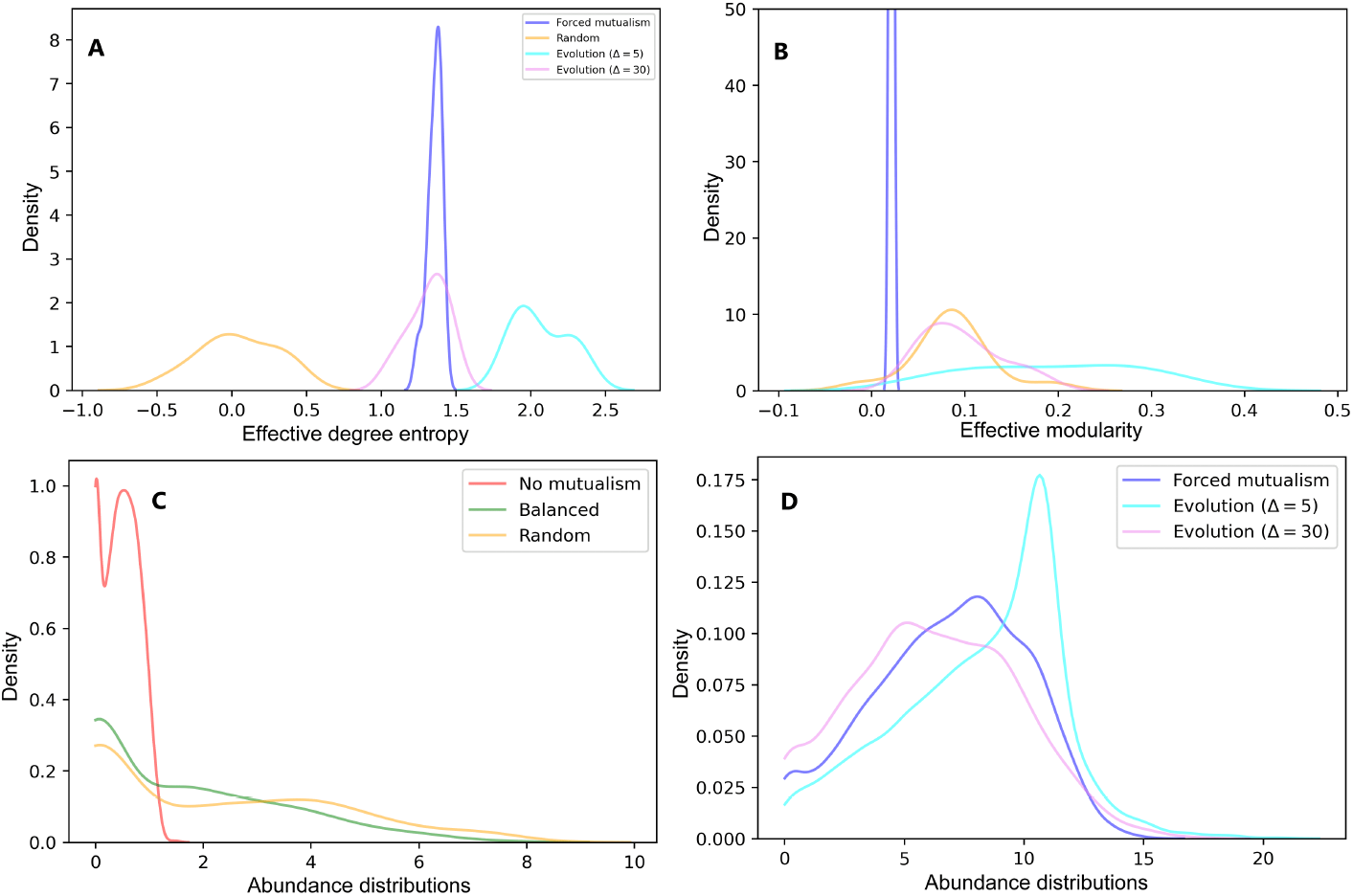
Complex communities have higher degree entropy, modularity, and species abundances. Mutualistic communities feature higher **(A)** degree entropy and **(B)** modularity than expected by chance. Both effects are stronger at lower connectance, as is the case of more controlled evolution (Δ = 5). We show effective values at the end of the simulation (see Methods). A value of zero corresponds to the level of a random network with the same richness and connectance. **(C)-(D)** The distribution of abundances is skewed to the right in complex mutualistic communities and skewed to the left otherwise. This effect is also more pronounced for more controlled evolution. We show kernel density estimations (KDEs) of histograms as representations of the corresponding probability distributions (see Methods).

**Fig. 7.**
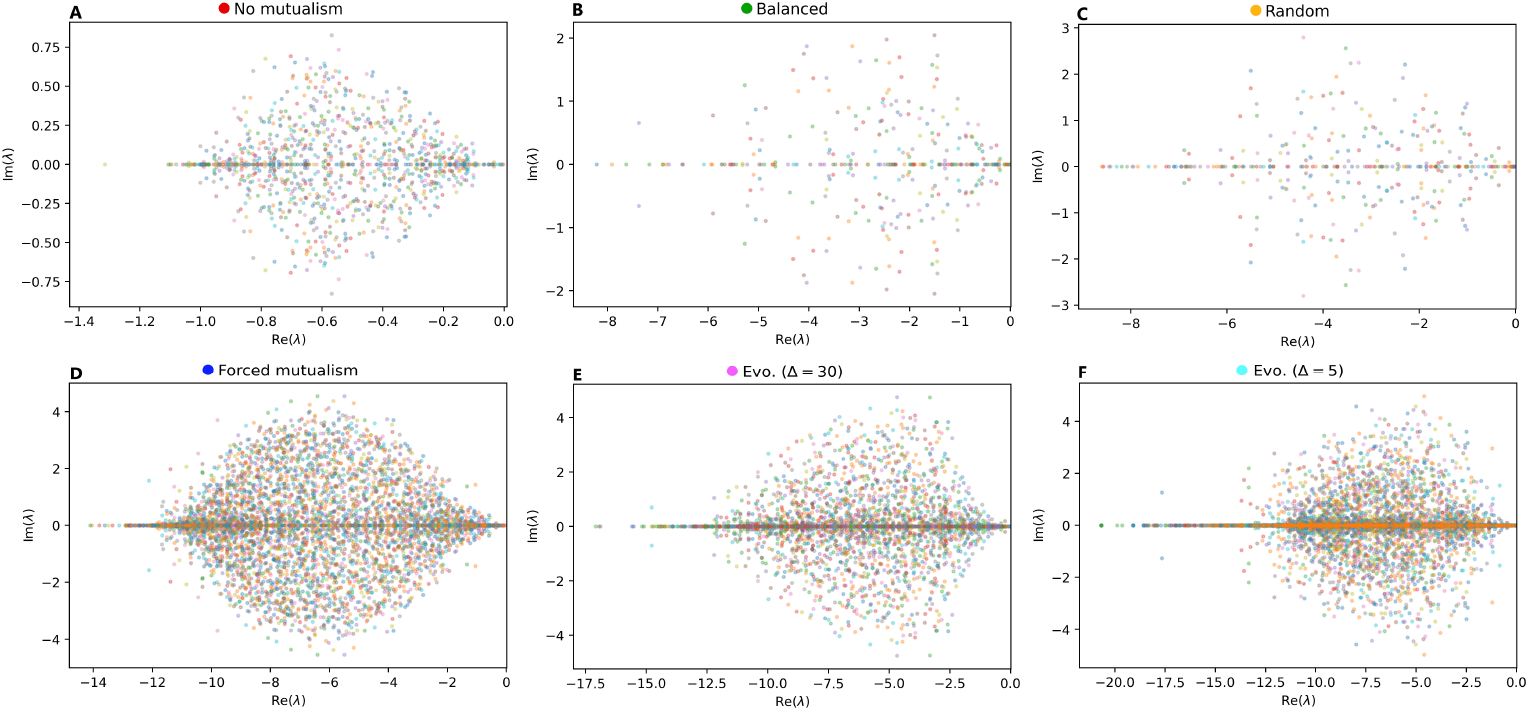
Evolution generates more stable networks. Distribution of eigenvalues, at the end of simulations, shown from lesser shifted to the left (A, invasion without mutualism) to most shifted (F, more controlled evolution). Points with different colours belong to different samples of assembled networks at the end of assembly trajectories. A high proportion of mutualism produces distributions of eigenvalues with a large shifts to the left. However, some evolutionarily assembled networks reach even further to the left than those from invasion with high mutualistic proportions. This longer line to the left, with points more concentrated on the real axes, is a product of evolution.

Another relevant feature of highly mutualistic communities is that, instead of a larger proportion of lower species abundances (i.e. rare species), it features a larger proportion of higher abundances (Figs 7C-D). This can have consequences for the balance of antagonistic interactions. In particular, communities without mutualisms exhibit more homogeneous distributions of lower abundances.

### 2.4 Evolved communities are more stable

Stability analysis of assembled communities shows a distribution of eigenvalues of the community matrix (see Methods) that is influenced by the proportion of mutualistic interactions. The distribution of eigenvalues is associated with stability against perturbations[26]. Larger proportions of mutualisms, both under ecological and evolutionary assembly, tend to shift the distributions to the “more stable” zone of the space (left-hand side in all panels of Fig. 7). Curiously, the forced mutualisms invasion model, which presents the highest proportion of mutualistic interactions, exhibits a slightly smaller shift than the evolutionary models. Moreover, evolution produces a concentration of eigenvalues on the real axis, which is associated with less oscillatory behavior of the system dynamics, and a shift to the left of the most negative eigenvalues that are concentrated along the axis. This suggests that mutualistic interactions emerging from evolutionary processes are more conducive of stability in complex communities than those emerging in ecological assembly models.

## 3 Discussion

The assembly of complex ecological communities is widely considered to be the result of evolutionary and ecological processes acting together to generate and maintain diversity. Yet, our knowledge on the ways in which both families of mechanisms can interact and produce this diversity is still limited. We have aimed at filling this gap through a theoretical investigation of the consequences of considering the evolution of species interaction networks within a dynamical community context, where the dynamics of species are governed by multiple interaction types. Our findings unveiled two fundamental results from simulations of community assembly by invasion or evolutionary speciation involving diverse interaction types. Firstly, we have found that highly mutualistic communities can grow to high levels of complexity while being robust to external perturbations. These predominantly mutualistic communities can form the foundations upon which food webs can also emerge through the accumulation of consumer-resource interactions. Secondly, evolutionary assembly via speciation emerged from our results as a key process to naturally generate those highly mutualistic communities. Depending on the amount of ecological variation between parent and offspring species, two types of complex communities can arise: communities with very high species richness or highly connected communities.

These two types of outcomes have been empirically observed in complex real ecosystems such as microbial communities. Furthermore, contrary to what has been previously suggested, we show that these outcomes do not require a limitation in complexity[27]. We thus argue that the apparent complexity-stability constraint on these communities may instead be the result of two different evolutionary settings depending on the adaptive traits of the microbes. Moreover, assembly by species invasion results in a strong convergence of outcomes, while evolutionary speciation can generate a variety of outcomes, ultimately producing a landscape of diverse complex communities. This aligns with the recognised importance of priority effects and historical contingency in the evolutionary assembly of microbial communities[28, 29].

Our result of a high proportion of mutualisms increasing stability while a low proportion having a destabilising effect on food webs agrees with previous results by Mougi and Kondoh (2012, 2014)[13, 30] and extend them to communities with another interaction type: interspecific competition. Furthermore, our results are more general as we do not rely on the assumption of constant effort across interactions. Our results thus confirm that these features of mutualistic communities do not come from constant effort of species across interactions, as previously suggested by Suweis et al (2014)[31]. The model presented here does not necessarily assume direct mutualistic relationships, but only reciprocal positive influences on species abundance, which could happen through the indirect environmental impact of species. High proportions of mutualisms enable the conditions for the coexistence of a diverse array of interaction types and an accentuated growth of the community. Conversely, a low proportion of mutualisms is not sufficient to maintain such conditions. We argue that the synergistic effects of widespread mutualistic interactions can promote the maintenance of a high abundance for the majority of species, enhancing persistence in the face of exploitative interactions. However, a small number of mutualistic interactions generates a high abundance of a small fraction of species, promoting perturbations that can tip off the balance of exploitative interactions. We suggest that a structural reason for this role of mutualisms is related to the way in which this structure affects species abundance distributions. The resulting shift of the distribution of eigenvalues to a more stable region might be a direct result of specific changes of these distributions[26]. Overall, our results agree with a positive complex-stability relation for mutualistic interactions and a negative complex-stability relation for antagonistic interactions[32].

Communities resulting from evolutionary assembly by speciation are in agreement with previous results showing that a path towards stability is reducing competition and increasing highly connected mutualistic interactions[16]. This is precisely the composition favored by selective pressures available in our models. However, this is not a definitive path, as it became evident how much diversity of outcomes an evolutionary process can generate. We suggest that a further specification of interaction types, along with constraints on mutualistic and consumer-resource interactions, could reveal other pathways towards a complex realistic community. Chomicki et al (2019)[15] reviewed the variety of roles that mutualisms can assume in impacting species richness, which can differ radically depending on their biological details and environmental conditions. We suggest that part of the plurality of the effects of mutualisms on species richness comes from its structural tendencies to increase complexity. This can be achieved in different ways, resulting in different outcomes for species richness, as we have shown. This effect is particularly dependent on the composition of interaction types.

Another possible path to evolve complex communities, shown here as a consequence of a lesser strength of species divergence from speciation, is to considerably increase species richness while maintaining a low connectivity between species in the community. This process produces highly rich and abundant communities that tend towards a larger modularity. In agreement with previous results on the relationship between mutualism and modularity[25, 33], this suggests that complex mutualistic communities can be selected for high modularity, and that network structural properties are sufficient for this emergence of more modular communities. The fact that an evolutionary mechanism is necessary for the natural emergence of complexity, diversity, richness, abundance, and modularity, is in agreement with previous predictions that networks should be sensitive to evolutionary constraints[20, 21].

Our results provide insights into the different processes by which complexity can arise during the assembly of ecological communities. The spread of mutualistic interactions and an evolutionary assembly by speciation have emerged as key mechanisms for these processes. Future work could expand on these results by exploring the consequences of evolutionary assembly in a meta-community context[34]. These efforts could focus, for example, on exploring whether dispersal between communities enhances the emergence of complexity and diversity and how to generalise the mechanisms we found for single communities. Additionally, the modeling of species traits and adaptive evolutionary dynamics[35] should inform relevant mechanisms generating patterns of specific biological communities, such as food webs[36–38] and microbial communities[39–41].

## 4 Methods

### 4.1 Model

#### 4.1.1 Ecological Dynamics

We define a generalised Lotka-Volterra dynamical network model of ecological community dynamics where species are represented as nodes and ecological interactions between them represented as weighted directed links. The model considers a mixture of interaction types by allowing for different combinations of sign and magnitude of interactions. Time-varying abundances are thus governed by the following set of differential equations:

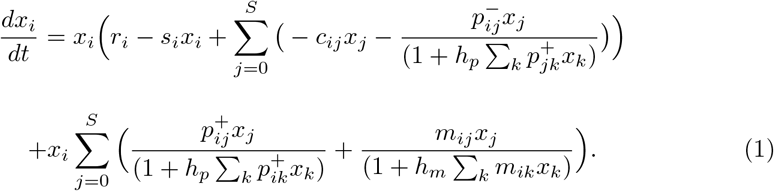

where *x*_*i*_ density of species *i* among *S* species in the community. Parameters *p*_*ij*_, *m*_*ij*_, and *c*_*ij*_ *>* 0 represent consumer-resource (+*/−*), mutualistic (+*/*+) and competitive (*−/−*) interactions respectively. These values determine the strength of the effect of species *j* on the density of species *i*. We thus consider three different types of interactions. For clarity, we represent 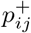 if *i* is a consumer and 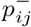 if *i* is a resource. Note that the effect of interacting species on one another is also proportional to their rate of encounter measured by *x*_*i*_*x*_*j*_.

The model assumes a Type II functional responses for saturation of intake of mutualists and consumers. For competition, there is no inherent saturation, and the resulting effect is governed solely by the encounter rate. Consumers and mutualists have their interactions saturated by handling and satiation times, and thus their capacity to grow is dampened by the amount of consumption or mutualistic benefits they acquire, respectively. Thus, within the functional response, *h*_*p*_ and *h*_*m*_ are the average saturation times for consuming and mutualistic interactions.

In addition to ecological interactions, species are subject to an independent reproduction rate *r*_*i*_ and intraspecific competition with strength *s*_*i*_.

#### 4.1.2 Evolutionary dynamics

Ecological dynamics as governed by Eq. 1 are altered by evolutionary events occurring at comparable timescales. Evolution in our model occurs via speciation with inheritance of interactions, thus adopting ecological relationships as a proxy for species traits. Speciation events occur once ecological dynamics have achieved equilibrium. Once the system governed by Eq. 1 is at ecological equilibrium (i.e. species abundances have attained long-term dynamical state), a ***parent*** species is randomly chosen from the community. A new ***offspring*** species is thus created which will ***inherit*** the ***parent’s*** interactions with modifications. Modifications are determined randomly with a strength of divergence Δ, defined as the maximum shift of interactions caused by removing inherited interactions and creating new ones.

In particular, whenever a new offspring arises, with interactions inherited from the ***parent*** species in the first instance, a ***magnitude of shift*** is randomly chosen from the integer interval [1, Δ]. Then, a random number of interactions are removed and created in a way that the sum of creations and removals amounts to the chosen shift. For example, if Δ = 5, a given offspring can have a shift from the parent of magnitude randomly chosen from 1, 2, 3, 4, 5. If the chosen value is e.g. 3, then one random possibility is to have 1 interaction removed and 2 interactions created, resulting in a shift of 3. This rule is versatile in terms of how much difference mutations can produce, and it also lets us fix how controlled the evolutionary process is. A high Δ means a loose evolutionary process, with less inheritability. Once the magnitude of shift is obtained, interactions from the offspring species to be removed, and interaction partners with which to establish new interactions are both randomly chosen. These are taken from the set of interactions of the offspring species and species in the community that are not currently interacting with the offspring species, respectively.

This ***magnitude of shift*** or ***mutation strength*** enforces an intrinsic trade-off between the conservation of information (smaller Δ) and more flexibility in the navigation of trait space (larger Δ). In our simulations, we consider two speciation modes, a more controlled one with Δ = 5, and a less controlled one with Δ = 30.

#### 4.1.3 Non-evolutionary assembly

To assess the extent to which complex species interaction networks emerging from the eco-evolutionary dynamics described above differ from purely ecologically assembled communities, we also implemented a non-evolutionary community assembly model grounded on invasion events.

As in the case of evolutionary assembly, invasion-driven assembly proceeds by adding new species to the community after ecological equilibrium has been attained. Differently from evolutionary assemble, the interactions of the invader are chosen from externally pre-defined rules, without the use of information from the community (i.e. no inheritance). Interactions of the invasive species are assigned with a probability that is randomly set between 0.2 and 0.7, which is its individual connectance. This allows for the connectivity of the community to adapt as species with different connectivities can emerge and be selected for by the ecological dynamics.

To test the effect of different interaction types on the final outcome of invasion-driven assembly we tested 4 different scenarios of invasion that impose different types of constraints on the invasive species:

1. **Random**. This is the baseline scenario in which the interactions of the invasive species are determined as explained above (i.e. randomly). This translates into the invader having any random proportion of interaction types.
2. **Balanced**. In this scenario the proportions of interaction types of the invasive species is constrained to always be balanced across types. Thus the invader is constrained to have 25% of each interaction type: mutualism, competition, consumer, and resource. This results in 50% of consumer-resource interactions. The total amount of interactions is still chosen as above (i.e. with a probability between 0.2 and 0.7 per species).
3. **No Mutualism**. New species are added to the community with balanced interactions (i.e. as scenario 2), but forbidding mutualistic interactions.
4. **Forced mutualism**. The proportion of mutualistic interactions in the invader is randomly chosen between 90% an 95%. This allows the simulation of highly mutualistic communities, which are not naturally emergent.

#### 4.1.4 Model simulations

Using the models described above we ran a series of numerical simulations to evaluate the effects of eco-evolutionary dynamics on network assembly and compare the resulting communities with communities assembled from pure ecological invasion dynamics. Each simulation proceeds by simulating ecological dynamics governed by Eq. 1 and introducing new species once ecological equilibrium is attained either via evolution or invasion (see scenarios above).

The assembly process starts with 5 non-interacting species for which initial abundances are drawn from the uniform distribution on the interval [0, 0.02]. An extinction threshold *x*_*ext*_ = 10^*−*5^ across all simulations. Whenever a species reaches the extinction threshold, it is removed from the system. Once a species are added into the ecological system of Eq. 1, it can cause the reorganisation of the community via e.g. species extinctions or changes in other species biomass. New species are included in the community with initial abundance that is equal to the extinction threshold *x*_*ext*_, and they are resampled until their growth from the threshold is positive. Once the system returns to equilibrium after each of these perturbation events, a new introduction event, either evolutionary or invasion-based occurs. We assembled networks until 1500 introduction attempts were reached. Therefore, if no species go extinct, our simulations would yield a maximum of 1505 species in the network.

To focus on the effects of interactions between species, we kept intrinsic growth and intraspecific competition rates, and interaction saturation times always constant, i.e. *r*_*i*_ = *s*_*i*_ = 1 and *h*_*m*_ = *h*_*p*_ = 0.1. Across all assembly scenarios, the strength of newly created interactions were drawn from a normal distribution 𝒩 (0, *σ*^2^), with the standard deviation *σ* = 0.5. The signs of the interactions was determined according to interaction type being instantiated following the assembly scenarios. To constrain consumer-resource interactions such that consumer biomass does not increase more than the reduction in resource biomass, we use an energy efficiency transfer fraction of 0.8, i.e.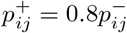.

We ran 15 simulations for the each of the scenarios above using the python library Scipy [42] for numerical integration.

### 4.2 Analysis

#### 4.2.1 Stability analysis

To perform linear stability analysis of the resulting communities, we numerically calculate the Jacobian matrix of the system from Eq. 1 at the equilibrium reached by the community after the occurrence of the last assembly event. The Jacobian is the matrix of derivatives of the right-hand side of Eq. 1, defined as 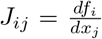, where 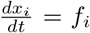. We obtain the eigenvalues *λ*’s of this Jacobian matrix and assess the stability of the system by representing the real and imaginary parts of the *λ*’s on a 2D plane.

#### 4.2.2 Network metrics

##### Complexity

We follow the classical measure of complexity in ecological networks established by May [9], 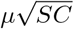, whereμ is the mean of all interaction strengths, *S* is the number of species and *C* = *L/S*^2^ is the connectance of the network (with *L* the total number of links). Complexity is also commonly defined using the variance of interaction strengths instead of the mean, although for random matrix models of zero mean without dynamical selection of weights[10]. May’s original work defines complexity in terms of the mean weights, as it directly reflects the intensity of ecological interactions. Considering *μ* as the mean or the variance of interaction strengths does not qualitatively change our results.

##### Entropy

The entropy of the network’s degree distribution *P* (*k*), calculated as *H* = *−*Σ_*k*_ *P* (*k*)*ln*(*P* (*k*)).

##### Modularity

A standard network metrics used to quantify the extent to which groups of nodes are more connected within them than to other groups[43]. It is based on nodes partitions (or sets of ***communities***) and given by

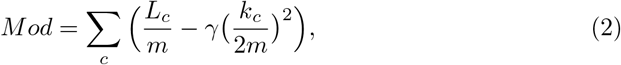

where *c* is the index of the community or module, *m* is the total number of edges, *L*_*c*_ is the number of intra-community links, and *k*_*c*_ is the sum of degrees of nodes in the community. The parameter *γ* is the resolution parameter, chosen to be 1. We implemented modularity using the python package NetworkX[44].

##### Effective measures

The network metrics described above are presented in what we define as effective values. We calculate them by subtracting the value obtained on the studied network for a given property from the value obtained for the corresponding Erdos-Renyi random network with the same size *S* and connectance *C*. Thus, the effective value of a metric *Z* is *Z*_*s*_ = *Z − Z*_*r*_, where *Z*_*r*_ is the value of the metric for the random network. In this way, we are interested in the shift from what would be expected by chance, representing a shift towards higher or lower values of *Z*.

We study the probability distributions of network metrics for different models built from histograms of values transformed into continuous kernel density estimations (KDE), for clearer visualization. For this, we use the Seaborn python package[45]. KDE is an algorithm aiming to produce smoother visualizations of histograms by estimating the underlying probability density function.

## 5 Statement of Authorship

GA and ML conceived and designed the study. GA implemented the model and performed numerical simulations and analysis. GA and ML wrote the manuscript.

## 6 Acknowledgements

This project was supported by the Leverhulme Trust through Research Project Grant # RPG-2022-114.

## Notes

### Competing Interest Statement

The authors have declared no competing interest.

### Summary of Updates

No significant differences with the previous version were introduced. Changes were mostly to writing style, reference list and manuscript formatting and the correction of typos.

